# Pan tissue analysis of human *NUMB* alternative splicing

**DOI:** 10.1101/2023.09.26.559629

**Authors:** Yangjing Zhang, C. Jane McGlade

## Abstract

Alternative splicing (AS) of pre-mRNA generates multiple protein isoforms from a single gene. It is developmentally regulated and mis-regulation is associated with many diseases. Previous studies that examined AS of the cell fate determinant *NUMB* demonstrated that it undergoes a regulated switch in the inclusion of protein coding exons 3 and 9 during rodent development and the differentiation of human cell lines. Here we extend this work by comparing exon 9 and exon 3 inclusion levels of the human *NUMB* gene across different normal human tissues by analysis of RNA sequencing data deposited in the Genotype-Tissue Expression (GTEx) and the Vertebrate Alternative Splicing and Transcription (VastDB) databases. Our results support earlier studies and reveal specific Numb isoform expression patterns in previously unexamined tissue and cell types, suggesting that Numb isoform expression is regulated in the normal development of a broad range of tissue types throughout the body.

## Introduction

Pre-mRNA undergoes splicing to remove introns and ligate exons to form mature mRNA. A single gene can generate multiple mRNA and protein isoforms by using alternative exons and splice sites, a process called alternative splicing (AS). AS occurs in nearly all multiexon genes and diversifies the transcriptome and proteome. The protein isoforms produced by AS can have different, or even opposite, biological functions and deregulated AS often plays major roles in disease including cancer (Park, Pan et al. 2018). The *NUMB* gene product is a cell fate determinant that undergoes AS of two coding cassette exons (exon 3 and exon 9) to produce 4 mRNA transcripts resulting in 4 protein isoforms: p72 (Ex9in, Ex3in), p71 (Ex9in, Ex3sk), p66 (Ex9sk, Ex3in) and p65 (Ex9sk, Ex3sk) (Dho, French et al. 1999) (Figure 1 describes alternative splicing of *NUMB* RNA and formation of the major NUMB protein isoforms).

**Figure 1.**
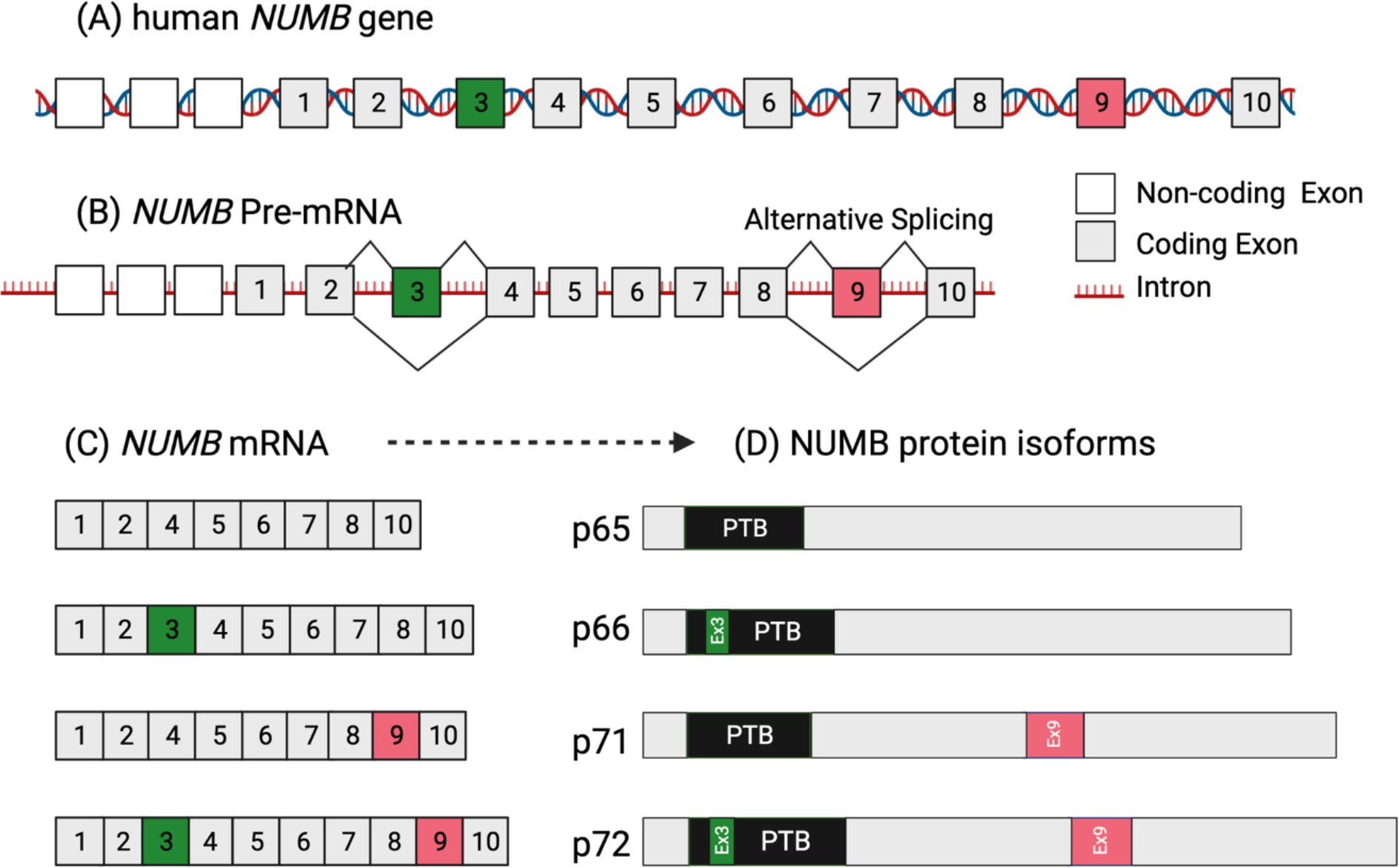
Formation of major NUMB protein isoforms by alternative splicing. (A) The human NUMB gene is comprised of 13 exons: 1-3 are non-coding and 4 - 13 are coding (ENSEMBLE transcript: ENST00000555238.6). Exon numbering used here is based on the coding exons. (B and C) Alternative splicing of exons in precursor mRNA forms the four major NUMB mRNA splice variants by inclusion or exclusion of coding exon 3 (Ex3in and Ex3sk respectively) and coding exon 9 (Ex9in and Ex9sk), and these are translated to NUMB protein isoforms identified here by their molecular weights: p65, p66, p71 and p72 (D).

Individual studies have documented the switch in splicing of *NUMB* exon 9 from inclusion to exclusion during development in rodents and differentiation of cell line models (Dho, French et al. 1999, Verdi, Bashirullah et al. 1999, Dooley, James et al. 2003, Corallini, Fera et al. 2006, Bani-Yaghoub, Kubu et al. 2007, Gao, Chi et al. 2011, Moran, Goldberg et al. 2011, Kim, Nam et al. 2013). Furthermore, reversal of this switch to re-express exon 9 is a frequent AS event in cancer (Chen, Chen et al. 2009, Langer, Sohler et al. 2010, Misquitta-Ali, Cheng et al. 2011). Comprehensive analysis of exon 9 and exon 3 inclusion levels of the human *NUMB* gene utilizing the accumulating RNA-seq data deposited in public databases has not been reported.

The Genotype-Tissue Expression (GTEx) database is a comprehensive public resource to study tissue-specific gene expression and regulation in normal human samples from approximately 960 donors. The Cancer Genome Atlas (TCGA) provides an opportunity for the in-depth analysis of the transcriptome and AS in tumors from over 8,700 patients with matched normal samples spanning 33 cancer types (Kahles, Lehmann et al. 2018). Our recent study of NUMB AS using these datasets (Zhang, Dho et al. 2022) confirmed and extended earlier work showing increased exon 9 inclusion in multiple cancer types. Here, we describe *NUMB* splicing of exon 3 and exon 9 across different normal human tissues, derived from the analysis of the GTEx RNA-seq dataset. In addition, we compare the results with that from VastDB, a database of AS profiles across multiple tissue and cell types with a more limited sample size (Tapial, Ha et al. 2017). Our analysis not only confirms, using a much bigger sample size, previous studies, but also reports additional human tissue types that undergo AS regulation in *NUMB*.

## Methods

### Data download

Alternative splicing analysis was conducted across RNA sequencing data from the GTEx database (Kahles, Lehmann et al. 2018). The PSI (Percent Spliced In index) values for exon 9 (chr14:73745988 - 73746132) and exon 3 (chr14:73783097 - 73783130) regions for *NUMB* and the raw read counts of all aligned genes was extracted from the supplementary tables together with sample IDs. The PSI values for exon 9 (HsaEX0044216) and exon 3 (HsaEX0044220) for *NUMB* and the cFPKM (reads per kilobase of target transcript sequence per million of total reads) of Numb and Numblike across different human tissues was downloaded from VastDB (http://vastdb.crg.eu/wiki/Main_Page) (Tapial, Ha et al. 2017).

### Expression analysis

GTEx samples were categorized into different tissue types using sample metadata supplied on the GTEx websites. A dotplot representing the mean PSI value for samples of each tissue type was generated for exon 9 and exon 3. The DESeq2 (version 1.24.0) R software package was used to normalize the readcounts for aligned genes from the GTEx datasets. A dotplot representing the mean normalized read counts was generated for each normal tissue sample from the GTEx database using R (version 3.6.3). Dotplots for PSI value of NUMB exon 9 and exon 3, and the cFPKM (reads per kilobase of target transcript sequence per million of total reads) value of Numb and Numblike extracted from VastDB, were also generated using R.

## Results and Discussion

The Percent Spliced In index (PSI) represents the percentage of gene specific mRNA transcripts which include a specific exon (Park, Pan et al. 2018) ^1^. We extracted the PSI value from the GTEx database for two cassette exons of the *NUMB* gene (coding exons 9 and exon 3) in different human tissues. There was an average of 52 samples per tissue type (Figure 1A, 1B, Supplementary Table 1) and the sample number, median/mean PSI value for NUMB exon 9 and exon 3, as well as total transcript level (in read counts) for NUMB are tabulated in Supplementary Table 1. Similarly, we extracted PSI values for *NUMB* exon 9 and exon 3 from the VastDB database, which includes AS profiles across multiple tissue and cell types but with a limited sample size (Supplementary Table 1) (∼2 samples per cell or tissue type) (Figure 1C, 1D). Together these data provide a comprehensive profile of *NUMB* isoform expression across 83 different human tissues or cell types at the transcript level.

A broad range in exon 3 and exon 9 inclusion level is apparent across different cell and tissue types. Tissues were grouped into different categories based on patterns of exon 9 and exon 3 inclusion level (Table 1). In human stem cells (ESC: embryonic stem cell and iPS: induced pluripotent stem cells), both exon 9 and exon 3 inclusion is greater than 50% suggesting that the p72 (Ex9in & Ex3in) Numb isoform is the predominantly expressed Numb isoform (Figure 1**C**).

**Table 1.**
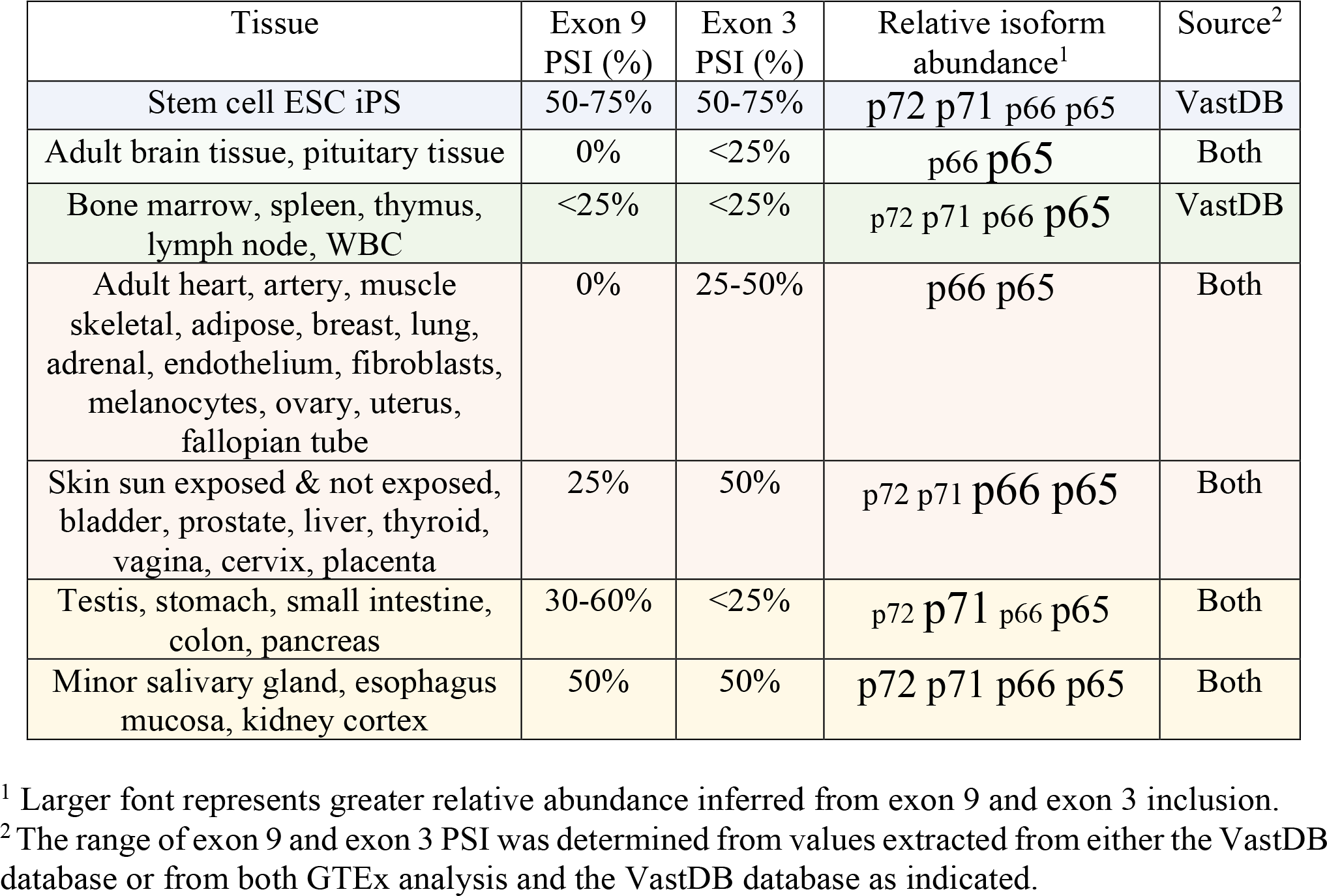
Numb isoform expression in normal human tissue.

In adult brain and pituitary gland, immune and hematopoietic system, on average both exon 9 and exon 3 inclusion level is lower than 25%, suggesting that the p65 (Ex9sk & Ex3sk) Numb isoform is predominantly expressed (Table 1). These observations are consistent with previously observed changes in both exon 9 and exon 3 inclusion during development measured using RT-PCR or protein expression: rat and human brain (Verdi, Bashirullah et al. 1999, Dooley, James et al. 2003, Bani-Yaghoub, Kubu et al. 2007), mouse pituitary gland (Moran, Goldberg et al. 2011), rat auditory epithelium (Gao, Chi et al. 2011), and human and mouse erythroid cells (Huang, Vu et al. 2021).

Heart, adipose, breast, lung, adrenal, and gynecologic tissue (ovary, uterus, and fallopian tube) have nearly 0% exon 9 PSI and 25-50% exon 3 PSI on average, and therefore are predicted to express the exon 9 excluded p66 (Ex3in) and p65 (Ex3sk) isoforms predominantly.

In tissues with self-renewal activity and stem cell populations such as the testis and gastrointestinal tissue (stomach, small intestine and colon, pancreas), the exon 3-excluded NUMB isoforms, p71 (Ex9in) and p65 (Ex9sk) are predicted to be predominantly expressed since they exhibit ∼50% exon 9 inclusion and less than 25% exon 3 inclusion. This is consistent with published studies which examined both exon 9 and exon 3 inclusion in the development of mouse pancreas (Yoshida, Tokunaga et al. 2003) and testis (Corallini, Fera et al. 2006). Other tissues such as salivary gland, esophagus mucosa, kidney cortex and prostate, have similar levels of included and excluded exon 9 and exon 3 transcripts therefore potentially express similar levels of all four protein isoforms.

The total transcript level of *NUMB* and its paralogue *NUMBL* were also compared across different human tissue types (Figure 2D). The mRNA transcript expression of *NUMB* appears to be lower in brain than in other tissues (Figure 2B). *NUMBL*, which exhibits some partially overlapping functions with *NUMB*, particularly in the nervous system (Petersen, Zou et al. 2002, Rasin, Gazula et al. 2007), is predominantly expressed in neural tissue. In most tissues, *NUMBL* is expressed at a lower level than *NUMB*, which is consistent with the literature (Figure 2D) (Zhong, Jiang et al. 1997, Gao, Chi et al. 2011).

**Figure 2.**
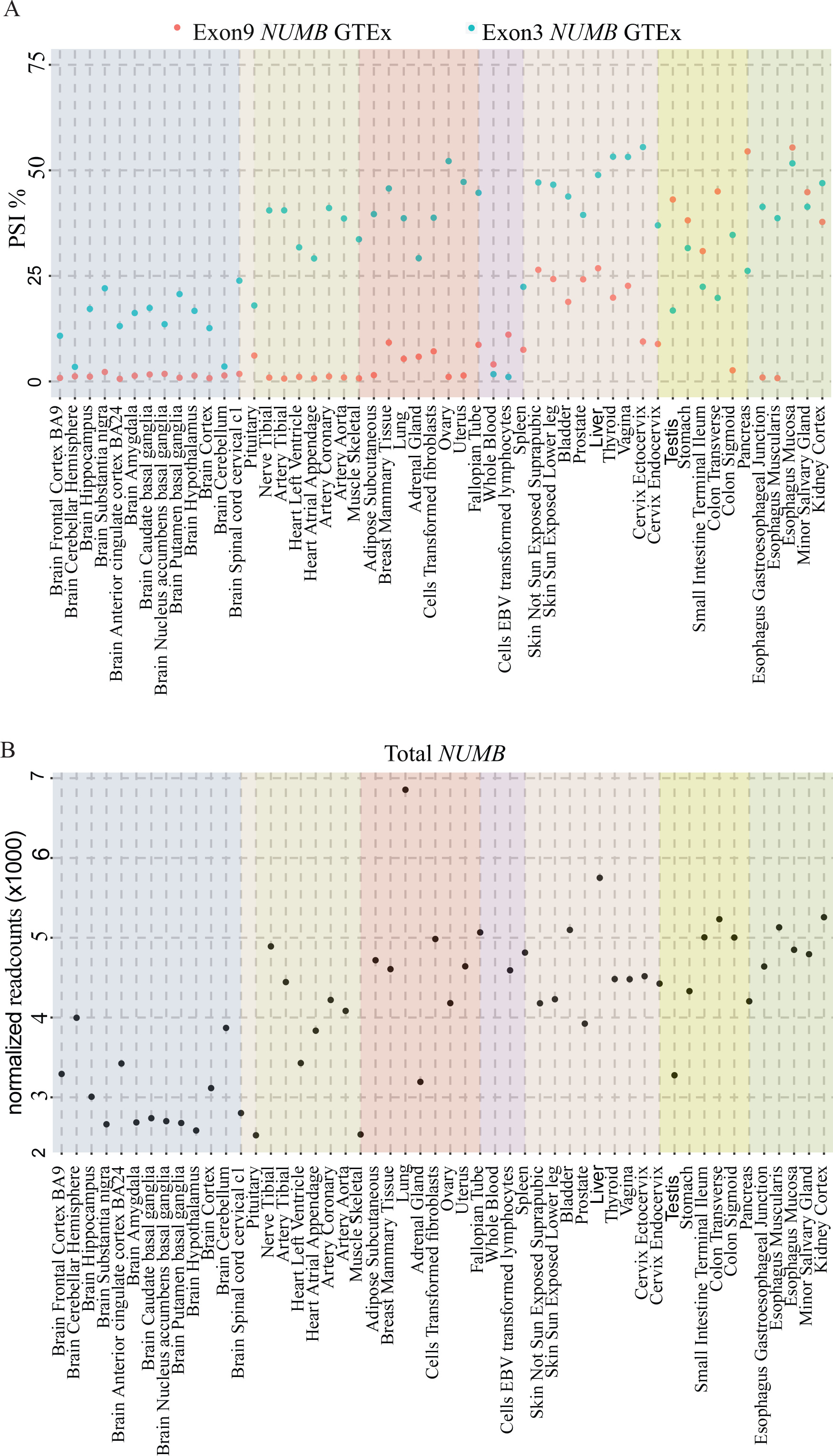
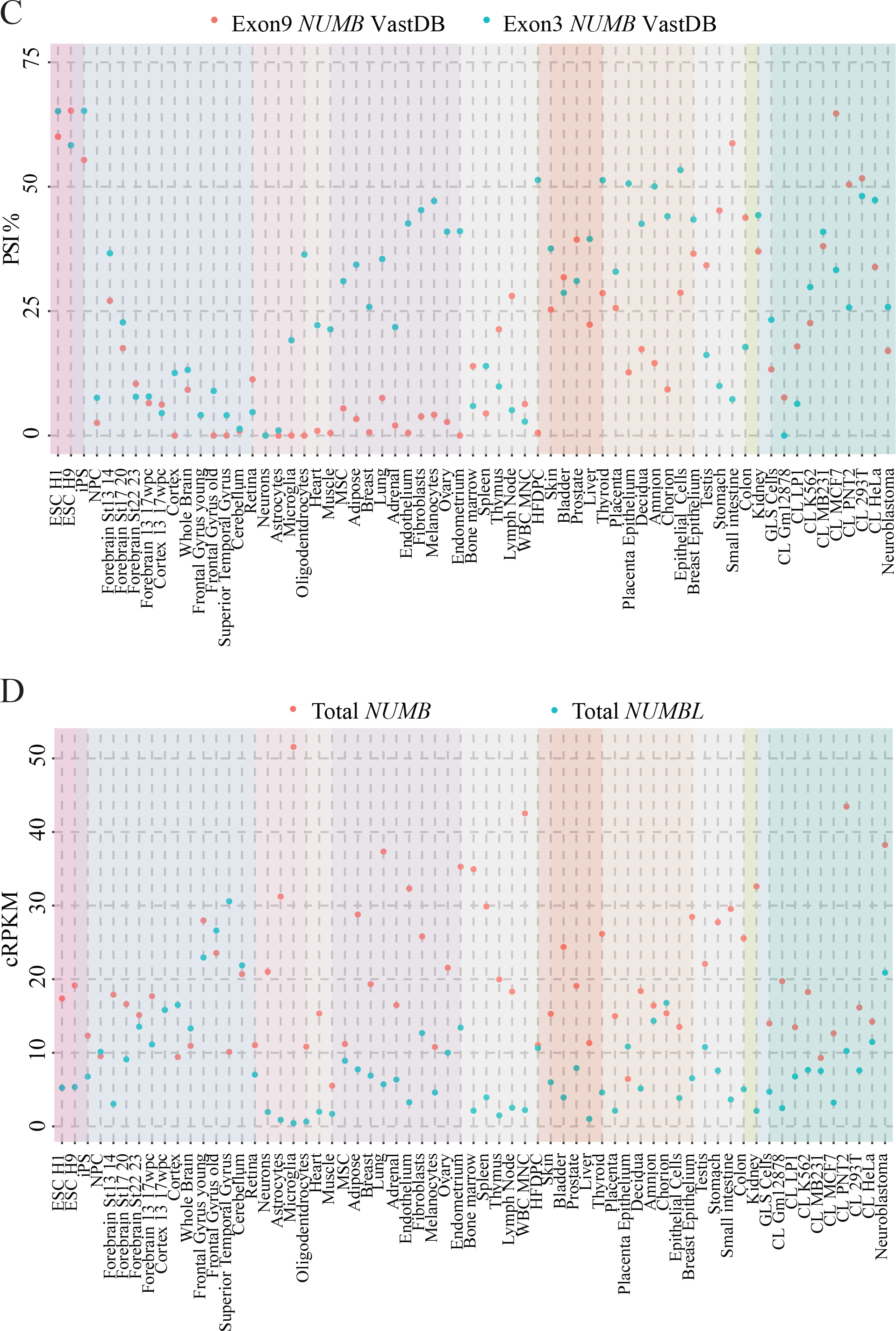
Mean inclusion level of exon 9 or exon 3 and total expression of *NUMB* (and *NUMBL*) in human tissues. **(A & C)** Exon 9 and exon 3 PSI in normal human tissues from the GTEx dataset (average sample size/tissue type = 52.3) (A) and in human tissues or cell lines from the VastDB dataset (average sample size/tissue type = 1.7) (C). Each dot represents the average PSI of samples for each tissue type. The represented tissues range from stem cells (ESC - embryonic stem cells, iPS - induced pluripotent stem cells etc.) to developing and adult neural tissues (different brain tissues etc.), glial cells (astrocytes, microglia, oligodendrocytes), muscle (heart, muscle), gynecologic tissue (ovary, uterus, and fallopian tube), genitourinary tissue (bladder, prostate, testis, kidney cortex), immune/hematopoiesis (bone marrow, spleen, thymus, lymph node, WBC), gastrointestinal tissues (stomach, small intestine and colon, pancreas) and other tissues. The rightmost columns for the VastDB dataset **(C)** are representative cell lines (CL). Primary neural progenitors, NPC. Hair follicle dermal papilla cells, HFDPC. Mesenchymal stem cells of adipose, MSC. Breast cancer cell line, MB231, MCF7. Prostatic cell line, PNT2. Lymphoblastoid cell line, Gm12878. Myeloma cell line, LP1. Chronic myelogenous leukemia cell line, K562. **(B)** Total transcript expression of Numb in different human tissues from the GTEx dataset represented as normalized read counts. **(D)** Total transcript expression of *NUMB* and *NUMBL* in different human tissue and cell lines from the VastDB dataset represented as cRPKM.

This analysis reports the tissue specific expression of *NUMB* exon 9 and exon 3 in human tissues and confirms isoform expression patterns previously identified in tissues from mouse and cell line studies. Importantly, our study also reveals the specific Numb isoform expression patterns in previously unexamined tissue and cell types including the heart, gynecologic tissues, genitourinary tissues, immune and hematopoietic system, suggesting that the regulation of Numb isoform expression is likely critical in a broad range of tissue types throughout the body and reflects different roles for the isoforms in development (Figure 2).

## Acknowledgements

The authors thank Dr. Sascha Dho for comments on the manuscript. This work was supported with funds from the Canadian Institutes of Health Research to CJM (FRN 106507).

## Author Contributions

YZ acquired, analyzed and interpreted data, drafted and critically reviewed the manuscript; CJM supervised YZ, analyzed and interpreted data, drafted and revised the manuscript.

